# *In silico* identification of non-coding RNAs in *Halobacterium salinarum NRC-1*, a model archeon organism

**DOI:** 10.1101/152439

**Authors:** M. A. S. Fonseca, R. Z. N. Vêncio

## Abstract

**Background:** In addition to the regulatory elements already known, for instance, transcription factors or post-translation modifications, there is growing interest in the regulatory role played by non-coding RNA molecules (ncRNA), whose functions are performed at a different level of biological information processing. Model organisms provide a convenient way of working in the laboratory, and different research groups use these models to conduct studies on the cellular mechanisms present in these organisms. Although some ncRNAs elements have been found in the *Halobacterium salinarum* model organism, we believe that not enough is known about these genomic regions.

**Methods:** Therefore, an *in silico* analysis for ncRNA identification was conducted on *H. salinarum* NRC-1. Considering a data integration perspective and some available methodologies, several machine learning models were built and used to designate candidate ncRNAs genome regions.

**Results:** A total of 42 new ncRNAs were identified. Combing analysis with other available tools, it had been observed that some suggested candidates also was found with different methodologies and thus, it highlights the proposed results.

## Introduction

Notably, the progress in biological knowledge has been widely guided by genomic data processing, where computational models emerge leading to a fuller understanding of biological mechanisms [8]. Model organisms have been used to discover general principles underlying more complex characteristics in all domains of life. Research based on the study of these organisms are oriented according to various interests, including those that are economical, agricultural and environmental, and those that involve human health [7]. The feasibility of model organisms for experimental studies is a great advantage, since they are easy to cultivate in the laboratory, and can be genetically modified [1, 7]. Among these, some research groups have worked with the archeal model organism *Halobaterium salinarum NRC-1* and several characterization analyses have contributed to our understanding of the organism and its use in industrial applications [17, 20]. Despite significant advances in previous studies related to *H. salinarum*, not enough is known about its non-coding RNAs (ncRNA) molecules. Is it know ncRNAs are involved in a wide range of biological processes, acting at different levels in the cell for information processing, including transcription regulation, replication, RNA modification and processing, mRNA stability and translation, and also protein degradation [19]. Due to its importance, many studies have been developed that aim to identify and characterize this class of molecules [16]. Computational approaches designed to identify ncRNAs have considered the inherent properties of such molecules, including sequence conservation and structure [14, 21], sequence length, transcript expression [10,12] and known functional motifs [2, 6]. Unfortunately, despite the existence of multiple methodologies to identify ncRNAs, it is difficult to rely on available strategies solely. Thus, in the present work, we developed an integrative *in silico* analysis to accurately predict new ncRNAs in *H. salinarum* NRC-1, aiming to contribute to the identification of these important regulatory elements. In order to ensure a significant strategy to select potential genome regions of ncRNAs, by complementing the available approaches, we also applied a Machine Learning (ML) based method to support our findings. Moreover, we gathered a collection of experimental data to increase the reliability of our results.

## Materials and Methods

### Currently available ncRNA prediction tools

Some conventional methods to predict ncRNAs are based mainly on primary sequence information. These approaches attempt to use homology and structure characteristics in order to perform their searches against ncRNA databases. The RNAspace platform http://www.rnaspace.org/ [3], for example, provides an integrated user friendly tool for ncRNA identification and annotation, whose methods are based on the mentioned characteristics. Other approaches exploit more specific experimental data as small RNA-seq (sRNA-seq) libraries; however, since these record mostly short length RNAs, is expect that particular features of ncRNAs be present in the sequenced reads. The approaches by both Dario [5] and Coral [12] try to use properties of sRNA-seq data based on mapped reads information. Another approach, named smyRNA [18], takes advantage of certain sequence motifs that are important in establishing the structure of the ncRNA molecule. These sequence motifs have a differential distribution across the genome and have exploited to identify new ncRNA region candidates. A further interesting methodology to identify ncRNAs involves the combination and integration of different data sources, since different properties capture distinct information about genomic elements [14].

### Data sources and training set definition

Available experimental strand specific data were used as relevant information for model building and further analysis. We integrated *H. salinarium sp*. NRC-1 data from small RNA-seq and tiling array data over 13 points from a standard growth curve [9]. We collected genome region annotations from the model organism, in order to represent previous knowledge, and making possible the training set examples collection for the machine learning techniques. Among these annotations, 2635 gene regions were obtained from [4]. Koide *et. al*. [9] reported 61 putative ncRNAs regions based on tiling array expression signal. Integrating several data types, they also identified 5’ and 3’ UTR, and we used this information in our approach. Additionally, we obtained 41 predicted ncRNAs from the UCSC Genome Browser (https://genome.ucsc.edu/). These candidates were raised from the snocan tool [13], which searches for motifs present in the C/D box snoRNA ncRNA class. The training set data corresponds to the available model organism annotated regions. As described above, we collected information about the genes (CDS), UTRs and already identified ncRNAs. In order to exploit these annotations and evaluate the predictive power of our ML model, we applied different training set configurations by manipulating the available genomic annotated regions. In one case, we used the full length of annotated regions considering the start and end of the original values. In another approach, we partitioned the regions according to [11]. All models were evaluated and the results will be described in the next section.

### ML features

Considering all genome annotated regions, we gathered available data sources corresponding to different categories and data information for the organism of interest (Table 1). The small RNA-seq signal corresponds to the counts of the aligned reads (in log 2 scale) and for each genome position a read-count value is associated, which indicates transcript expression. Since the transcripts are fragmented and diverse, the read count signal becomes unclear with several breaks, decays and oscillations. To improve the signal representation, aiming to handle this signal diversity, we tried to consider the read-count shape with some ML features, including kurtosis, skewness, mean, median, standard variation, interval (max – min values) and percentage of expression above the mean of the all read-count in the region.

**Table 1.**
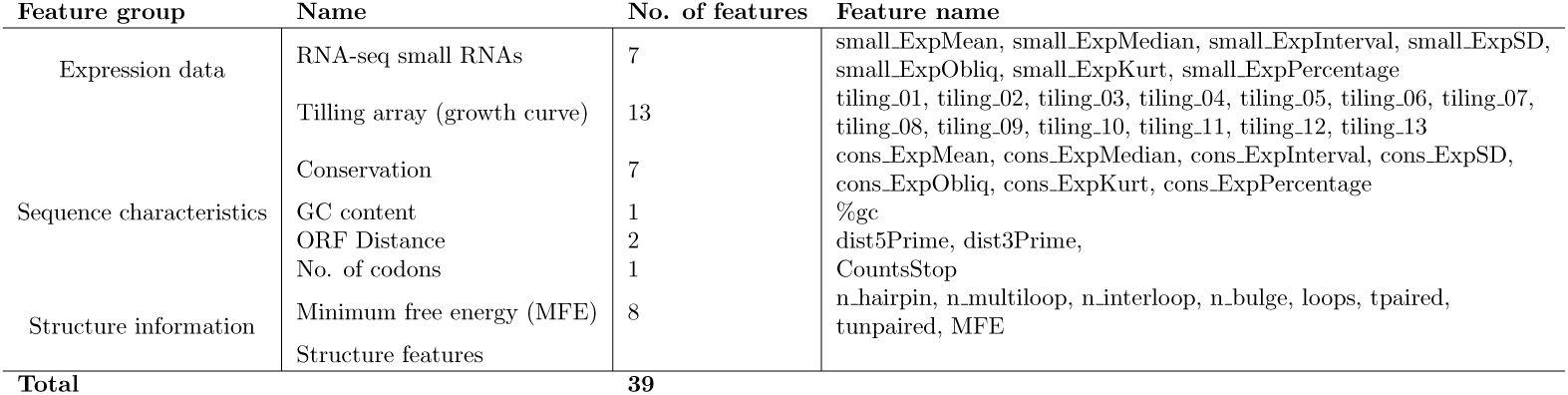
Summary of the machine learning features.

Another relevant information that helped us to distinguish the classes is the codon triplet sequence. We considered all start and stop codon definitions from an Archaea genetic code table, then we calculated the nucleotide distance from the interested region to the closest start and stop codon. This was called the open reading frame (ORF) distance feature. The sequence conservation measure was based on a previous method, as described in Marchais *et. al*. [15]. Considering a BLAST hit (https://blast.ncbi.nlm.nih.gov/Blast.cgi) for each genome position, the conservation index indicates the number of genomes on each position, and was weighted by its phylogenetic proximity with the *H. salinarum* NRC-1 genome. To handle the conservation information, we considered the same measures described above (kurtosis, skewness, etc), which was applied to the small RNA-Seq data. We also included GC content as part of sequence characteristics. Finally, secondary structure information was included based on the Context Fold tool prediction results (https://www.cs.bgu.ac.il/negevcb/contextfold/) [22]. The structure prediction annotation was then parsed and the sub-structures were obtained as a collection of features. In summary, 39 features were used (Table 1).

### ML model evaluation and statistical analyses

To precisely evaluate the predictive power of the ML classification model, several standard performance measures were used, including accuracy, sensitivity, specificity, ROC analysis and area under curve (AUC). These statistical evaluations involved the analysis of model hypothesis variance and bias, estimated from independent test sets outside of the training sets, and the cross-validation technique provided this assessment. Non conventional measures were also considered to evaluate the ML model. Since many biological data systems usually appear noisy, conventional classification measures may not properly reflect the model behavior. Thus, we applied a new strategy based on sliding window fragments to evaluate the prediction behavior sensibility.

## Results

### Identification framework for ncRNAs

To identify new ncRNA genomic regions candidates, we have combined both a newly developed ML methodology and available tools to predict ncRNAs. The main procedures of the developed approach are illustrated in Figure 1. First, the input data were processed in order to define ML features, using both available genomic annotation and representative information over these regions, such as experimental expression data and sequence properties (conservation, predicted structure). Considering the ML model, a sliding window strategy was applied across the entire genome. In general terms, the strategy splits the genome into several overlapping fragments, then uses these fragments as inference for the ML model. Subsequently, the probability ncRNA signal is obtained by manipulating the probability associated with each fragment. We defined peaks of high probability using signal processing procedures and then considered overlapping peaks to define candidate ncRNA regions. Finally, the final candidate regions were evaluated and filtered considering different experimental data and methodologies.

**Figure 1.**
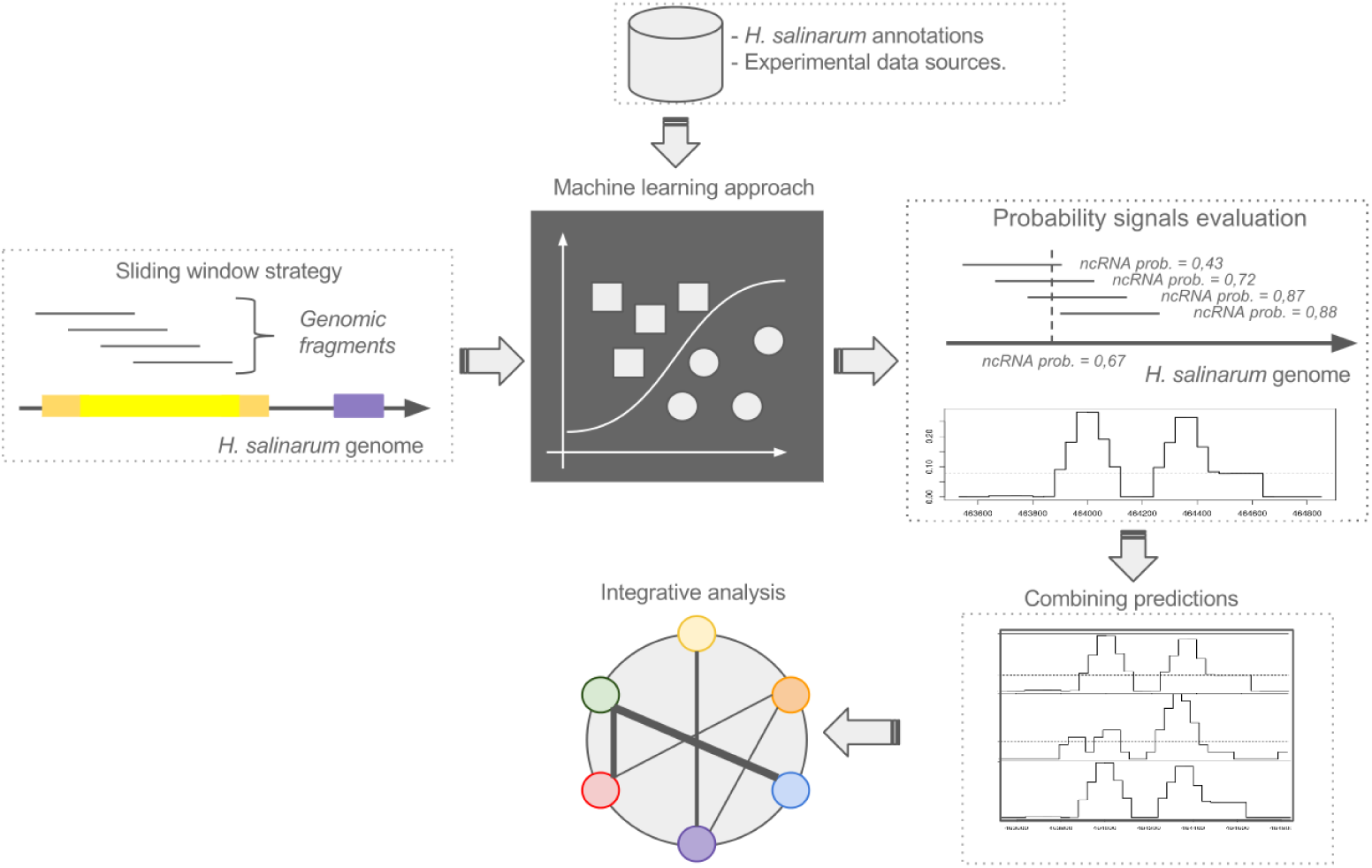
Workflow of the applied non-coding RNA prediction.

### Predictive behavior of the ML model

To ensure an unbiased evaluation of our ML models, we used a procedure that involves a sliding window prediction strategy across the entire *H. salinarum* NRC-1 genome. This procedure helped us to investigate the predictive behavior of the developed ML models, whose modifications reflect the different scenarios that we considered, by manipulating the genomic annotations of *H. salinarum* NRC-1, which are used as training sets. The first model (M1) uses the original information regarding the values of start and end of each annotated region (coding sequence, CDS; untranslated region, UTR) and the already known ncRNA available from [9]. In the second scenario (M2), each annotated regions (CDS, UTR) were fragmented, considering a fixed size of 120 nt [11]. To visualize the performance of the applied algorithms, using these two different training sets, the area under the curve (AUC) was plotted in Figure 2.

**Figure 2.**
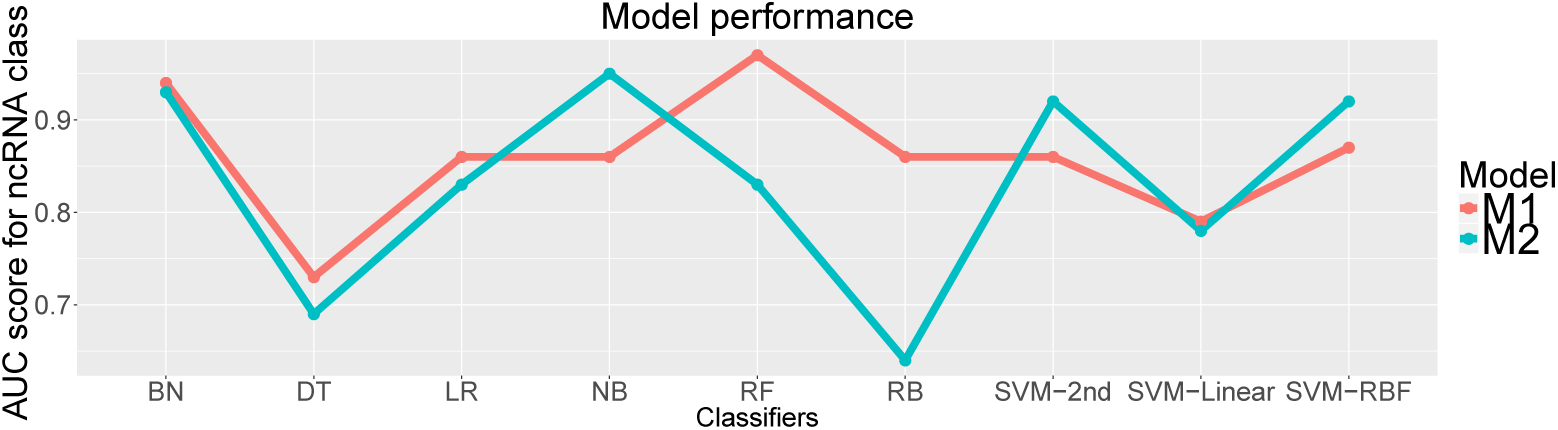
Area under the curve score showing the performances of the classifiers in 10- fold cross validation. We considered nine classifiers (Bayes Net, BN; Decision Tree, DT; Logistic Regression, LG; Naive Bayes, NB; Random Forest, RF; Rules Based, RB; and support vector machines (SVM) with tree kernels - polynomial, linear and radial basis function (RBF)). Two models, M1 and M2, were generated manipulating the training set annotations.

Among the nine algorithms tested, random forest (RF) achieved the highest AUC (of 0.97) with 10-fold cross-validation experiment and training set without fragmentation (M1), which suggested a good separation of the training classes. However, no clear elucidation of the predictive model behavior on unannotated regions arised from these results. To assess the overall classification sensitivity, we applied the sliding window strategy, considering the top 3 classifiers, based on the AUC measure in both models (Figure2). In total, 50354 fragments were used in this step, which covered all bases of the chromosome at plus strand. The fragmentation followed the same considerations described in [11]. After conducting inference process for each fragment, we found the ncRNA class probability value assigned to each genomic position. To map the overlapping fragments cases, all overlapping positions were taken together by the mean. We identified the most important regions (with high ncRNA probability) using a segmentation signal approach, which basically defined the start and end of each peak by checking the probability value variation, by comparing each position with the mean of all probability signals. To precisely evaluate the ML model prediction sensibility, we have compared the annotated regions, also used as training set, with all segmented peaks obtained earlier. Peaks clearly matching the CDS or UTR regions were counted as false positives.

A summary of the total peaks obtained is shown in Figure 3. According to the results, Bayes Net algorithm on M1 had 44.8% peak overlap annotation (CDS, UTR and ncRNA classes); 44.5% of them were false positive peaks. RF and support vector machines-radial basis function have 41% and 31% of peaks overlapping annotations, respectively. The total number of generated peaks also suggests an unwanted model prediction behavior, since a large amount of peaks are more favorable to increase the false positive results. For instance, the RF approach produced 2232 high ncRNA probability peaks. Based on these considerations we have noted that M2 achieved a better prediction behavior results, including relatively few total peaks and a reduced number of overlapping annotations. The Bayes Net classifier had, for example, 8.5% peaks matching with CDS regions and 21.3 % matching with UTR regions. Interestingly, the majority of false positives corresponded to UTR class (Figure 3). In summary, our results show that when we partitioned training regions (M2), the signals peaks displayed a more distinctive signature: reduced number of high ncRNA class probability candidates and few of them mapped to annotated regions. In order to better visualize these features, we plotted, using Gaggle Genome Browser (http://gaggle.systemsbiology.net/docs/geese/genomebrowser/), the probability signal over the entire chromosome of *H. salinarum* considering plus strand (Figure 4). Indeed, the high peaks are clearly distinctively across the whole genome range and are mainly located in intergenic regions.

**Figure 3.**
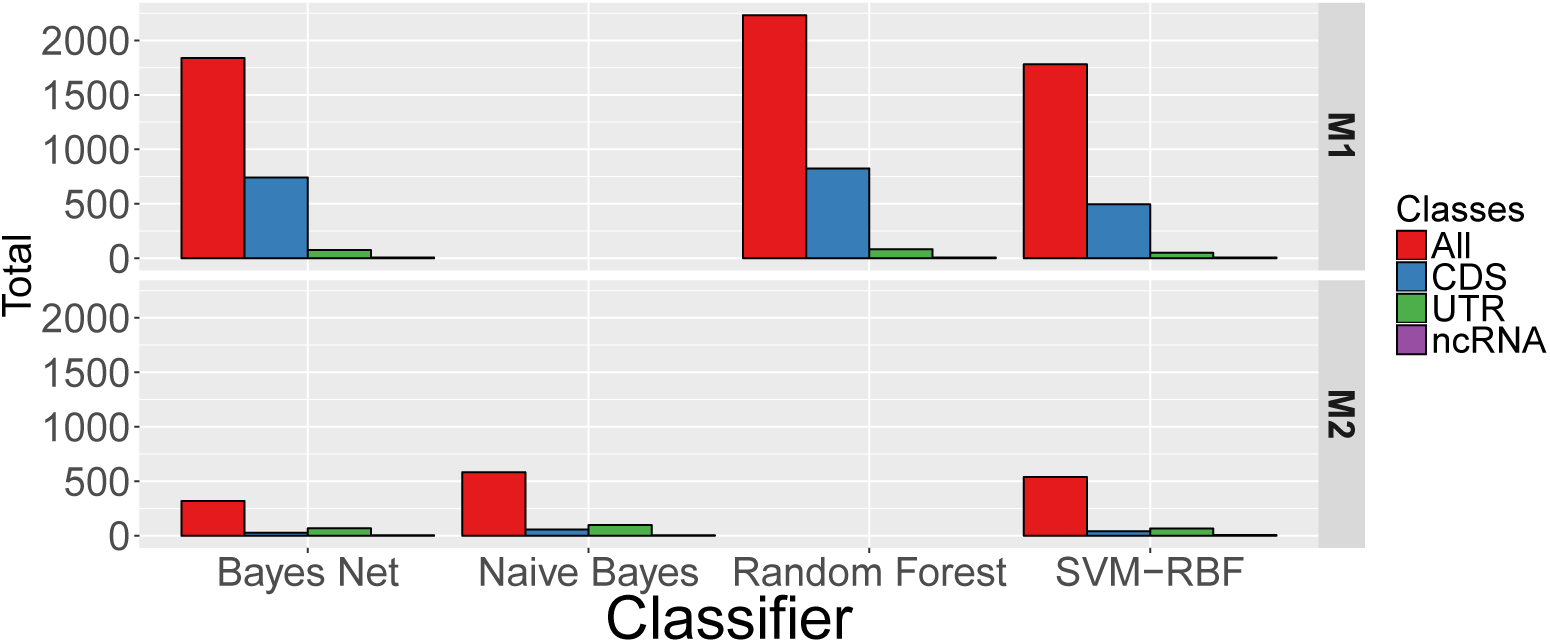
Summary of peaks obtained by each classifier. The red bars indicate the total peaks obtained in the top 3 area under the curve classifiers. We crossed the peaks against all training annotations (CDS, UTR and ncRNAS). Peaks that overlap CDS regions were considered as false positive.

**Figure 4.**
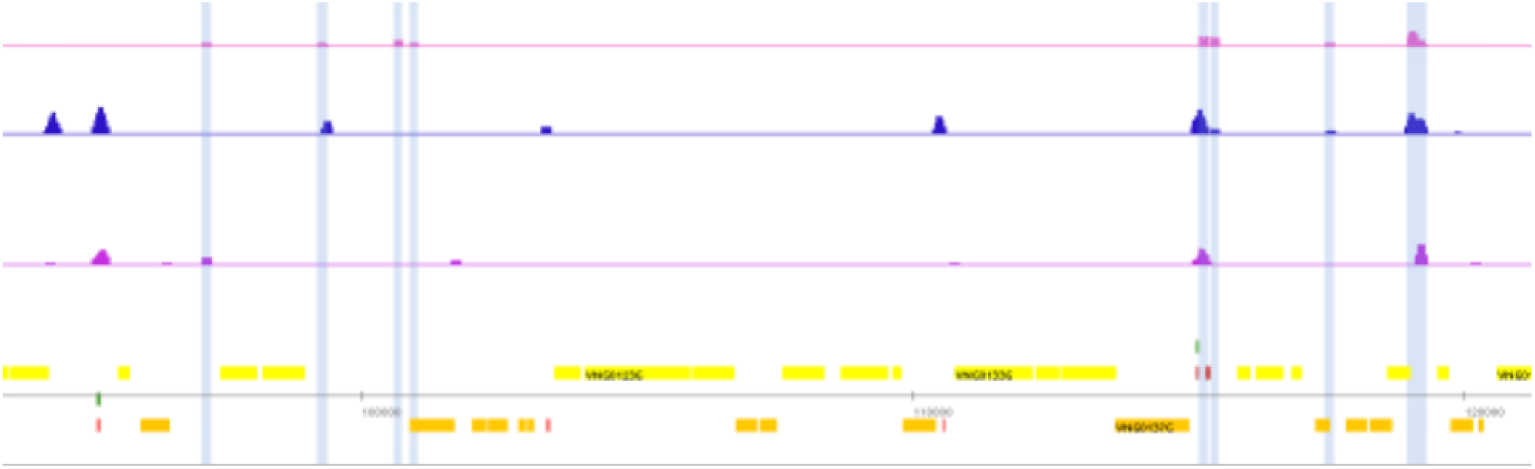
Example of peaks over the genome. From Gaggle Genome Browser window we can see tree tracks representing the probability of ncRNA signals in all genomic positions. The highlighted probability peaks in pink was obtained from naive Bayes results considering the M2 scenario. Boxes in yellow and orange indicates gene annotations in the forward and reverse strand, respectively

### Combing classification peaks

The prediction peak results obtained by each classifier independently were integrated using voting systems - regions greater than 400bp were removed. We compared all peaks and selected those that intersected the same region of at least five classifiers. By varying the number of overlapping predicted regions threshold value, the number of candidates can be increased. On the other hand, it creates the risk of many false positives. Here, we have opted to select at least 8 classifiers to chromosomes and 7 to plasmids, pNRC100 and pNRC200. After this selection step, some combined candidate regions were removed by the following criteria: clear false negatives (regions covering 90% of CDS regions and UTR), true positives (regions matching [9], ncRNAs annotations) and regions overlapping known tRNA or rRNAs. As a final result, 162 filtered regions emerged for further inspection.

### Candidates regions selection criteria

Aiming to include a better characterization of all 162 ncRNAs candidate regions and consequently to improve the evidence of a truly positive ncRNA class, we applied a visual and manual inspection using the Gaggle Genome Browser tool. In addition, for expression data, used as ML features, we also considered the RNA expression profile during the *H. salinarum* growth curve. Moreover, we used tiling array probe intensities for reference conditions for *H. salinarum* [6] and the relative enrichment of the aligned start position, from the primary transcript library available in [23]. Based on this new experimental information, we filtered the candidates according to the following criteria: absence or weak signal of the aligned start position enrichment (Figure 5A), weak tiling array peak signal (Figure 5B), regions that followed the CDS expression profile behavior (Figure5C), and regions with short overlapping positions (Figure5D). At first, we discarded candidate regions that were close to CDS coordinates, since it was hard to distinguish between UTR and genuine ncRNA classes. However, when we subsequently compared the 162 initial candidates with Transcription Start Site Associated RNAs (TSSaRNAs data, available in Zaramela *et. al*. 2014 [23]), we surprisingly noted that 40 regions overlapped the same TSSaRNAs regions results. This was interesting, since both methodologies are distinct and converge, in some cases, to the same findings. TSSaRNA-VNG1213C (Figure 6) was experimentally evaluated in [23]. The peaks defined by the classifiers were clearly high in the ncRNA genomic region. Both growth curve expression profile and reference wild type condition expression show changes across the highlighted area. There is an enrichment of aligned start reads overlapping the 5’ region. Based on the mentioned considerations we manually inspected all 162 initial candidates and produced 42 new regions as *H. salinarum* ncRNAs candidates. Some of these are differentially expressed in the growth curve. Moreover, all candidates have shown an enrichment of starting read information.

**Figure 5.**
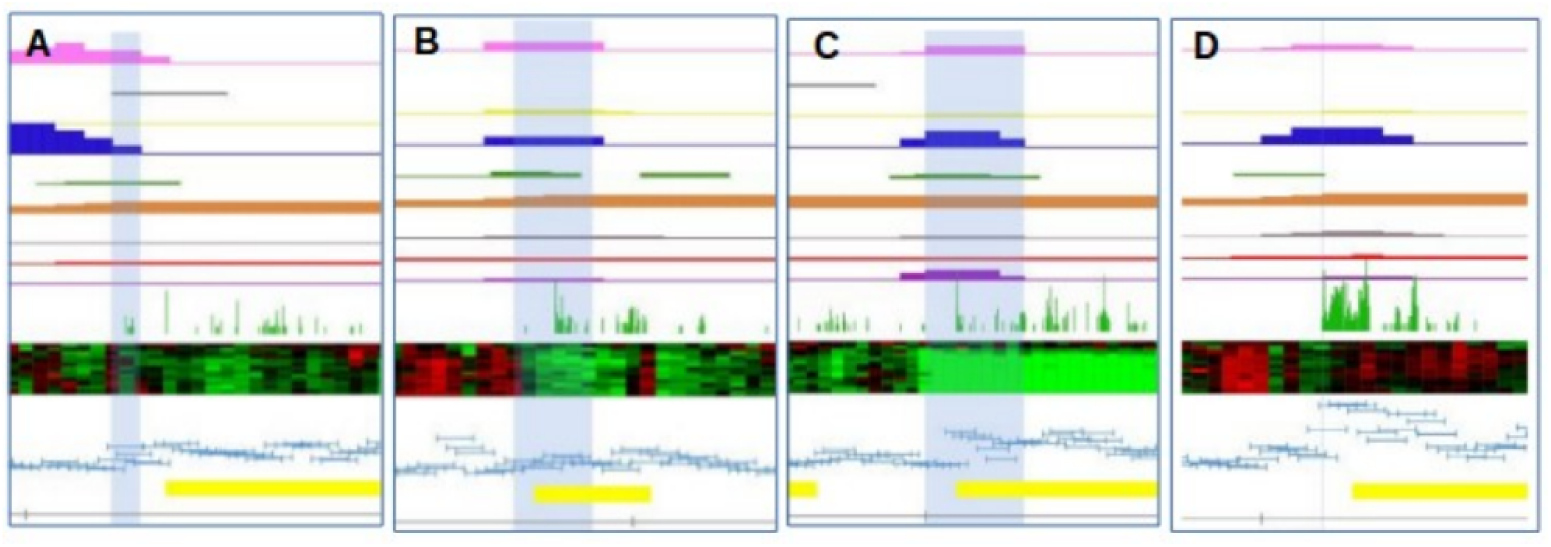
Examples of removed candidates. The highlighted regions indicate the evaluated region and the criteria in consideration. The first nine tracks corresponds to probability of ncRNA signals obtained by each classifier. The green bars corresponds to start read enrichment. The heatmap corresponds to 13 expression signals of *H. salinarum* in the growth curve. Finally, the light blue track corresponds to tiling array signals in the reference condition.

**Figure 6.**
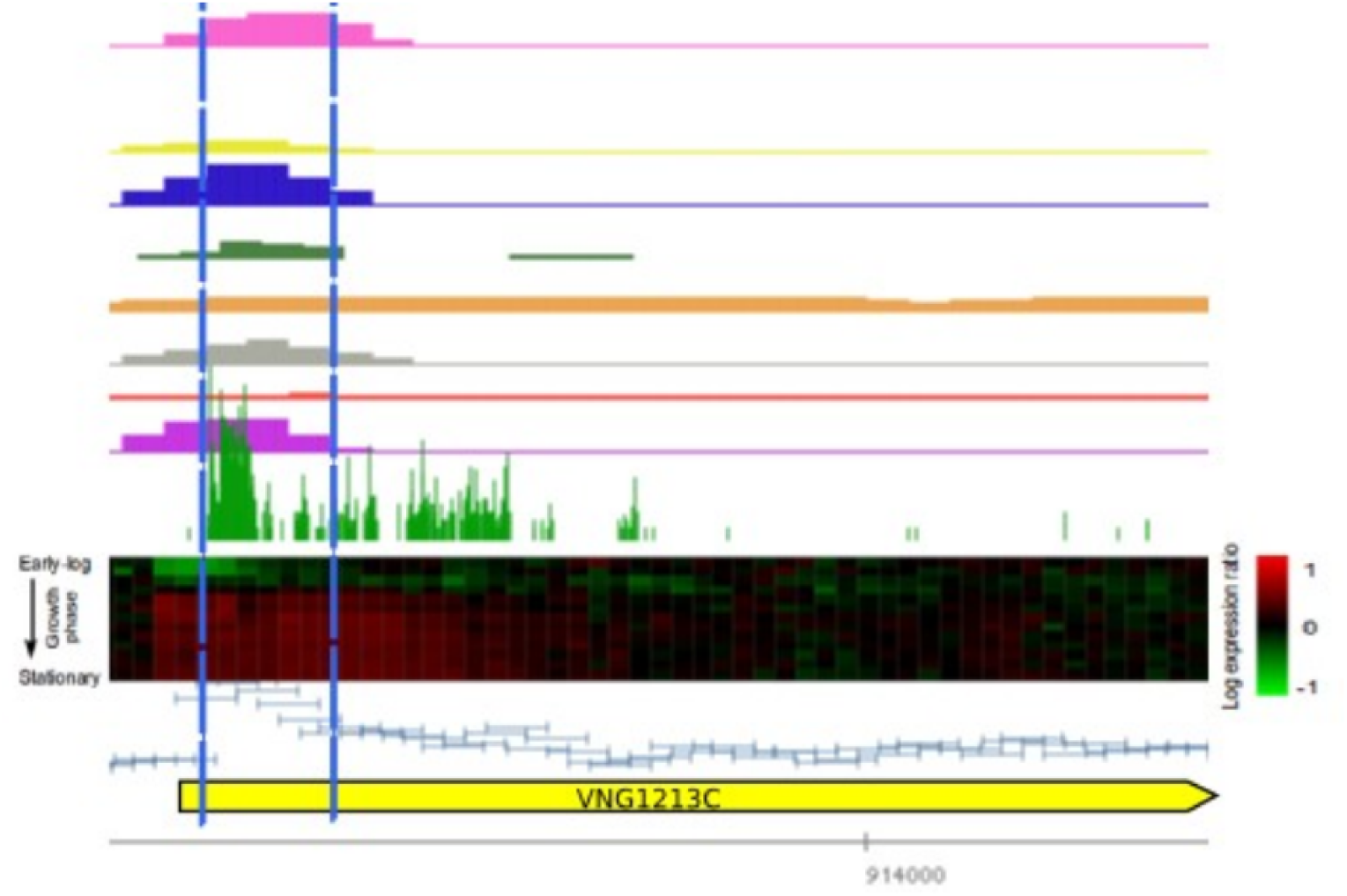
TSSaRNA-VNG1213C was experimentally evaluated in [23]. The peaks defined by the classifiers were clearly high in the ncRNA genomic region. Both growth curve expression profile and reference wild type condition expression show changes across the highlighted area. There is an enrichment of aligned starting reads overlapping the 5’ region.

### Integrative prediction results

For a further assessment of the newly identified ncRNA genomic regions, we integrated predictions combining results of different methodologies. We applied some available tools and summarized as follows: similarity based approaches (YASS and BLAST), essentially identifying regions related to tRNAs and rRNAs. The tRNAscan-SE tool also identifies tRNAs, since it applies a more specific search. The majority of known tRNAs and rRNAs also were identified by Darn (https://carlit.toulouse.inra.fr/Darn/index.php) and ERPIN (https://bioinformatics.ca/links directory/tool/9822/erpin) tools. The Darn approach also suggested 5 snoRNAs-CD-box overlapping: VNG1529G, VNG1726G, VNG0318G, VNG1585Cm and VNG1988G genes. These results were not confirmed by Snoscan [13], since they did not match with snoRNAs available in the UCSC Genome Browser. ERPIN found two regions related to small nucleolar RNA, overlapping VNG1654G and VNG2176H genes. RNAz (https://www.tbi.univie.ac.at/wash/RNAz/) obtained more interesting results, only two predicted regions overlapped annotated CDS. We observed that 22 regions correspond to UTR and 26 with annotated tRNAs. INFERNAL (http://eddylab.org/infernal/), RNAmmer (http://www.cbs.dtu.dk/services/RNAmmer/) and AtypicalGC tools predicted few, not clearly defined, regions. RNAmmer only found rRNA, and INFERNAL identified the RNaseP annotated as VNGs01; 9 regions predicted by INFERNAL and 14 obtained using AtypicalGC overlapped CDS. In summary, considering all prediction results, about 90% of the tools successful identified regions belonging to tRNAs and rRNAs. Since many regions predicted as ncRNAs in fact overlapped CDS annotations, we observed many false positives and subsequently, we had difficulty in evaluating the prediction results independently. Therefore, we opted to report just regions filtered by the developed ML approach, highlighting those candidates that were predicted under at least one of applied approaches. In total, 17 candidates (in bold) of 42 matches with other approach results (Table 2).

**Table 2.**
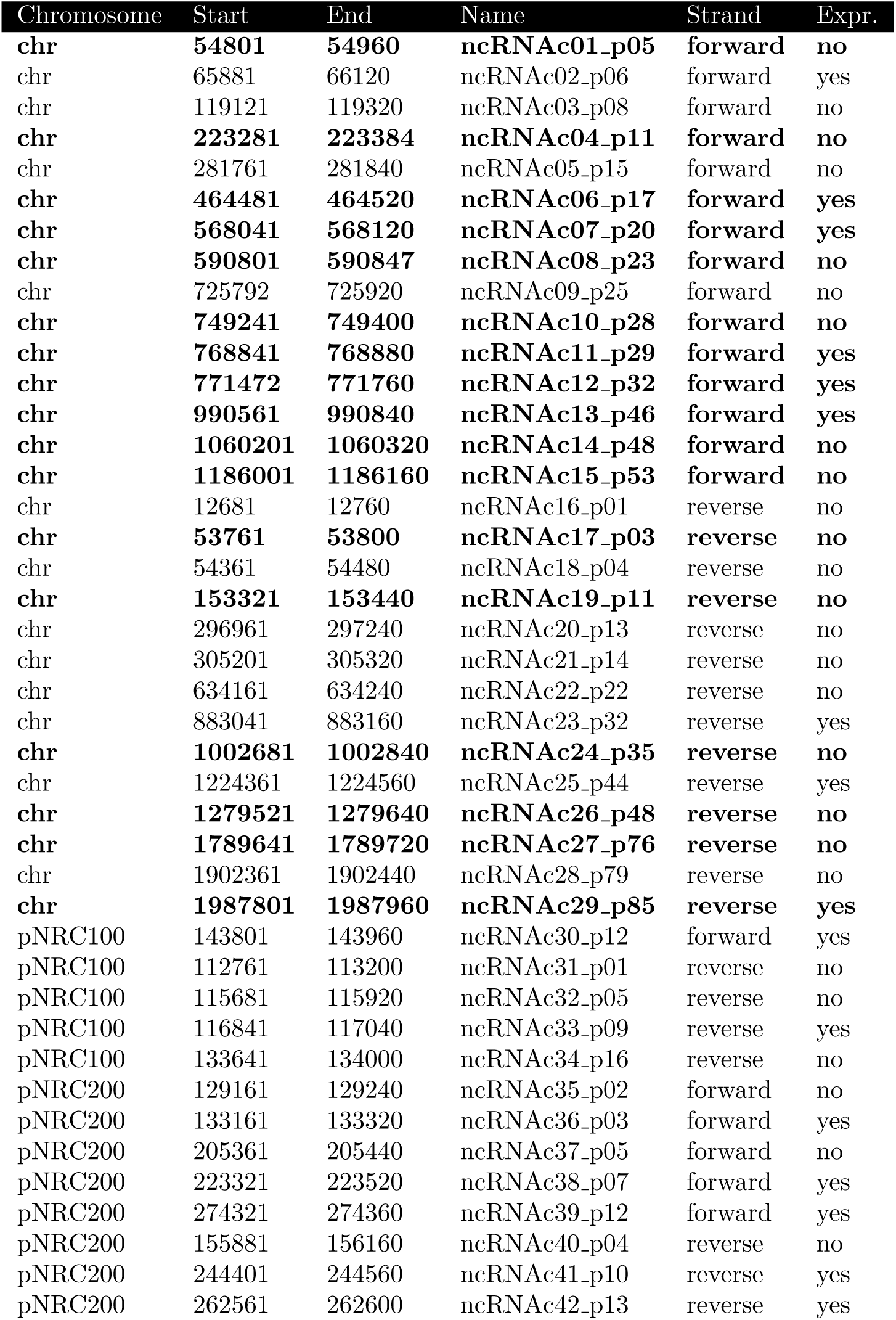
Genomic regions of identified non-coding RNA candidates. Candiates that were also identified by at least one of the applied approaches are highlighted in bold. The *Expr* column indicates if the region has expression variation along the growth curve.

## Discussion

Model organisms offer a convenient and extensive way for research. Different research groups aiming to guide their studies for a mutual and wide understanding of the cellular mechanisms present on these organisms. The transcriptome complexity includes not only translated transcripts, but a diversity of functional elements. The regulation of gene expression occurs at several cellular levels, and are also guided by non-coding elements. Identification of these non-coding molecules is a challenging task. Although some ncRNAs elements have been found in the *Halobacterium salinarum* model organism, we believe that not enough is known about these genomic regions. Therefore, we applied an *in silico* analysis for ncRNA identification to the *H. salinarum* NRC-1 genome. Considering a data integration perspective and some available methodologies, several Machine Learning models were built and used to designate candidate ncRNA genomic regions. We summarize our whole list of novel ncRNA candidates suggested by this work in Table 2. In total, 42 new regions were suggested as ncRNAs in *H. salinarum* NRC-1. The sliding window approach achieved the most significant results, overcoming traditional ML performance measures. Available methodologies were applied and helped to find more evidence in the final results. We had difficulties in evaluating candidate regions near to CDS, since it can also be associated to UTR regions; however, we compared 162 candidate regions with Zaramela *et. al*. [23] TSSaRNA results, and 25% of them were also found using a distinct methodology and offer support to our findings. We believe that the final work-flow can be automated and applied to other organisms (allowing comparisons with other approaches).

## Author contributions

All the authors contributed to the drafting of this manuscript.

## Competing interests

No competing interests were disclosed

## Grant information

MASF received scholarships from Fundação de Amparo à Pesquisa do Estado de São Paulo (FAPESP 2012/02896-9) and CAPES/Brazil.

## Acknowledgments

We are grateful to Diego Martinez Salvanha and Tie Koide for their kind help in providing important discussions and insights.

